# Biophysical properties of human β-cardiac myosin with converter mutations that cause hypertrophic cardiomyopathy

**DOI:** 10.1101/065649

**Authors:** Masataka Kawana, Saswata S Sarkar, Shirley Sutton, Kathleen M Ruppel, James Spudich

## Abstract

Hypertrophic cardiomyopathy (HCM) affects 1 in 500 individuals and is an important cause of arrhythmias and heart failure. Clinically, HCM is characterized as causing hyper-contractility, and therapies are aimed toward controlling the hyperactive physiology. β-cardiac myosin comprises ~40 percent of genetic mutations associated with HCM and the converter domain of myosin is a hot spot for HCM-causing mutations, but the underlying primary effects of these mutations on myosin's biomechanical function remain elusive. We hypothesize that these mutations affect the biomechanical properties of myosin, such as increasing its intrinsic force and/or its duty ratio and therefore the ensemble force of the sarcomere. Using recombinant human β-cardiac myosin, we characterize the molecular effects of three severe HCM-causing converter domain mutations R719W, R723G and G741R. Contrary to our hypothesis, the intrinsic forces of R719W and R723G mutant myosins are decreased compared to wild type, and unchanged for G741R. Actin and regulated thin filament gliding velocities are ~15 percent faster for R719W and R723G myosin, while there is no change in velocity for G741R. ATPase activities and the load-dependent velocity change profiles of all three mutant proteins are very similar to wild type. These results indicate that the net biomechanical properties of human β-cardiac myosin carrying these converter domain mutations are very similar to wild type or even slightly hypo-contractile, leading us to consider an alternative mechanism for the clinically observed hyper-contractility. Future work includes how these mutations affect protein interactions within the sarcomere that increase the availability of myosin heads participating in force production.

One Sentence Summary: Biophysical analysis of human β -cardiac myosin with converter domain HCM mutations reveal minimal changes compared to wild type myosin to slightly hypo-contractile profiles in ensemble force production, in contrast to the clinically observed hyper-contractility, leading to a speculative alternative mechanism involving availability of myosin heads participating in the sarcomere.

## Introduction

Hypertrophic cardiomyopathy (HCM) is the most frequently occurring inherited cardiac disease, affecting more than 1 in 500 individuals *(1)*, and 10% of HCM patients develop fatal arrhythmia and/or heart failure *(2)*. It is characterized by left ventricular hypertrophy in the absence of predisposing conditions (e.g. aortic stenosis, hypertension), ultimately causing decreased left ventricular chamber volume and obstructive physiology during systole. In 1990, Geisterfer-Lowrance et al. *(3)* reported a missense mutation, R403Q, in the β-cardiac myosin heavy chain gene (MYH7) in a cohort of HCM patients, and since then hundreds of different mutations in not only myosin but also other sarcomeric proteins (e.g., myosin binding protein-C (MyBP-C), troponin I and cardiac actin) have been identified. It is estimated that missense mutations in the β-cardiac myosin heavy chain (MyHC) are responsible for ~40% of cases of genotype-positive HCM *(4, 5)*. The principle pathology is manifested at the level of the ventricle in HCM *(6)*, and the conventional view is that HCM mutations result in hyper-dynamic cardiovascular physiology that is often seen as a supra-normal ejection fraction (EF) on echocardiograms *(7, 8)*. However, the molecular mechanisms that determine how missense mutations in the human β-cardiac MyHC cause the hyper-contractile phenotype remain unclear. In particular, while there has been more understanding of secondary cellular events that occur during the hypertrophic process, the primary effect of the mutations on the function of the sarcomeric contractile proteins is largely unknown. Hence, there is a paucity of the knowledge that is essential for development of new therapeutic treatments for this deadly disease, as well as for predicting the phenotypic outcomes in this heterogeneous patient population.

In order to understand the primary effect of HCM mutations, we are focusing on the molecular parameters of the contractile properties at the sarcomere level and are measuring the effect of HCM-causing mutations using in vitro reconstituted systems. These parameters include: 1) the intrinsic force (F_intrinsic_) of myosin, which can be increased or decreased by mutations that change the spring constant of the elastic element of the motor; 2) the duty ratio of the myosin, which is the fraction of myosin bound to actin in a force-producing state in the sarcomere at any moment during systole, estimated from the time myosin spends strongly bound to actin (termed t_s_) divided by the total actin-activated myosin ATPase cycle time (t_c_). The duty ratio can be changed in more than one way, such as a change in the ADP release rate (which determines t_s_) or the
weak-to-strong transition time (which determines t_c_); and 3) the velocity of actin displacement by an ensemble of myosin molecules, which is related to the stroke size (d) divided by the time myosin spends on actin (t_s_). Finally, the force produced by a given sarcomere can be expressed as the ensemble force (F_ensemble_), which is the effective force produced by a given sarcomere, expressed as a function of F_intrinsic_, the duty ratio of the myosin, and the total number of functionally available myosin heads overlapping with the actin in the sarcomere (N_a_): F_ensemble_ = F_intrinsic_ (t_s_ / t_c_) (N_a_). The experimental setups
for measuring these parameters have been rigorously explored previously *(9-11)*. Our overall hypothesis has been that HCM-causing mutations increase one or more of these parameters and thereby increase the power output of the cardiac muscle, which is the F_ensemble_ times the velocity of contraction. We *(10-13)* and others *(14, 15)* have proposed that such changes serve as the initial trigger for the pathologic hypertrophic process.

Early studies using reconstituted systems used mouse α-cardiac myosin as a model system. Those studies showed considerable increases in either ATPase activity, velocity of actin filament gliding in in vitro motility assays, or F_intrinsic_ *(14, 15)*, and hyper-contractility was easily explained by these significant increases in these fundamental parameters. However, more than 80 residues are different between mouse α-cardiac and human β-cardiac myosin; it has been reported that the same HCM mutation (R403Q) caused different mechanochemical effects on cardiac myosin depending on the myosin isoform in mouse *(16)*, and there is a significant functional difference between human α-cardiac and β-cardiac myosin (*11*). Thus it is important to examine the effects of the HCM mutations in the proper backbone, by using the human β-cardiac myosin. Recent studies using human β-cardiac myosin carrying the R453C mutation *(10)* or the R403Q mutation *(12)* failed to show such large changes in the same parameters, and whether the net result of the changes observed were contributing to a hyper-contractile or hypo-contractile state was difficult to assess.

We therefore sought to examine these fundamental parameters for human β-cardiac myosins carrying mutations in the converter region of the molecule, which is a hotspot for HCM mutagenesis *(17)*. To date, every variant known in the converter region in the human population results in HCM pathogenesis *(17)*, which is not true of any other region on the myosin molecule. Generally, HCM-causing mutations in the converter have been reported to result in worse clinical outcomes, while a wide range of phenotypes have been observed *(18, 19)*. Prior studies using mammalian and Drosophila muscle myosin showed that the converter is involved in determining the stiffness of the cross-bridge bound to the actin filament *(20)*, and therefore the intrinsic force generating ability of the head. Converter movement is coupled to ATP hydrolysis, phosphate (Pi) release and the force-generating event *(21, 22)*. Furthermore, converter movement is coupled to load-dependent ADP release, and thus ADP affinity could be changed, which would affect the maximum velocity of the motor, the duty ratio, and hence the net F_ensemble_ *(21-24)*. Thus, the converter is pivotal in transducing the energy of ATP hydrolysis to the stroke of the lever arm and force production and one might suspect that F_intrinsic_ or one of the other fundamental parameters would show significant increases compared to wild type. Strikingly, this is not what we have found, as shown in this report, and our results suggest that we need to rethink the molecular basis for clinically observed hyper-contractility caused by myosin-based HCM mutations.

Three HCM mutations were selected for this study: R719W is a mutation found in the early days of HCM genetics and has been characterized to be one of the most lethal mutations in MYH7, the gene encoding the b-cardiac myosin heavy chain *(25)*. R723G is a mutation first reported in families in Barcelona, who manifested progressive heart failure as well as sudden death *(26)*. G741R is also one of the longest recognized mutations found in multiple families *(27)*. Prior studies have shown that slow skeletal muscle fibers from soleus, which use the same human β-cardiac myosin heavy chain as the heart, albeit different light chains, containing the R719W mutation from HCM patients showed increased isometric force, increased stiffness in rigor and increased ATPase activity, but no change in kinetics of active cross-bridge cycling *(20, 28)*. A mouse model harboring the R719W mutation in a-cardiac myosin, generated by the Seidman group *(29)*, was shown to have activation of proliferative and profibrotic signals in non-myocyte cells that promotes pathological remodeling leading to HCM *(29)*. The R723G mutation has been studied using human skeletal muscle biopsy samples and showed similar changes, while there was no change in ATPase activity *(28)*. These mutations are also known to cause variable calcium sensitivity *(30)*. A recent study by Kraft et al. *(31)* showed that in mechanically isolated human cardiomyocytes from biopsy samples with the R723G mutation, maximum force was significantly lower and Ca^2+-^sensitivity was unchanged. Conversely, R723G mutant soleus fibers showed significantly higher maximum force and reduced Ca^2+^-sensitivity compared to controls. These discrepancies were speculated to be due to changes in protein phosphorylation in different muscle types *(31)*. G741R myosin was also studied from human soleus fibers and showed decreased velocity and force production *(32)*, whereas a study using a mouse myoblast cell line showed increased tetanic fusion frequency without significant impact on the contractile kinetics *(33)*. These previous studies suggest that there is a significant impact on the myosin function when the converter domain residues are altered *(21-24, 34)*, although conclusions are variable and complicated by differences in light chain composition, isoform being studied, post-translational modifications, disease state of the tissue used, and so forth. To date, there has been no report studying the effect of converter domain mutations on biomechanical properties of myosin function using purified human β-cardiac myosin. Here we report the primary effects of these mutations in the converter domain of human β-cardiac myosin on fundamental parameters of power output using homogeneous, expressed and purified human β-cardiac short S1, the motor domain of myosin. Our approach enables the direct comparison of wild type and HCM mutant forms of human β-cardiac myosin and eliminates the background variability inherent in using patient-derived samples that may contain non-synonymous variants *(35)*.

## Results

### The intrinsic force of R719W and R723G human β-cardiac short S1 is slightly decreased, while G741R shows no change compared to wild type myosin

For all of our experiments, we used the purified short subfragment 1 (sS1) construct of human β-cardiac myosin from residues 1 to 808 (Fig. 1A, Fig. S1A), which contains the catalytic domain and a short lever arm with only the human β-cardiac myosin essential light chain (ELC), and is known to be the motor domain of the molecule *(36-38)*. The same construct was used previously in other mutational studies of HCM *(10-12)*.

**Figure 1.**
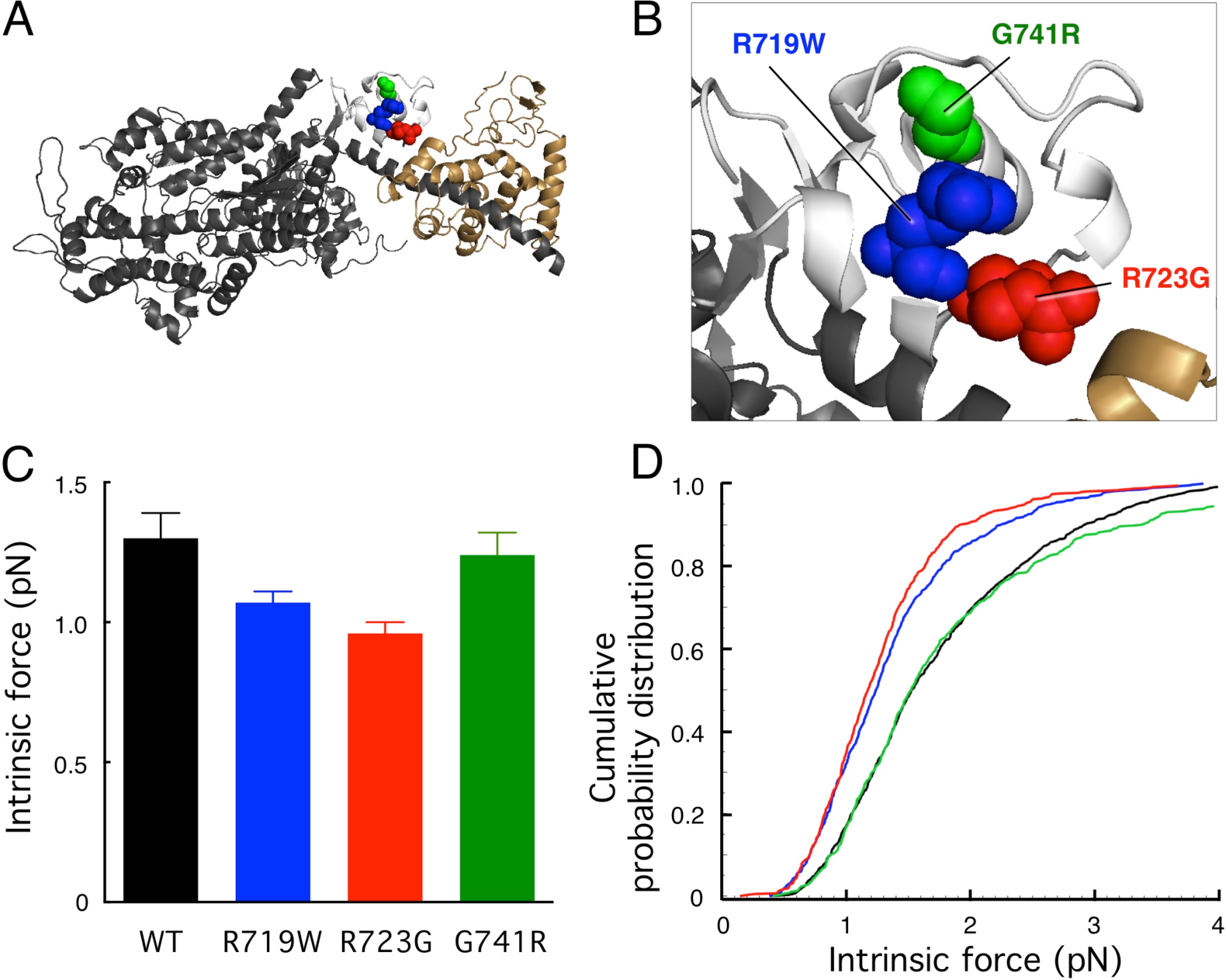
Structure of a homology-modeled human β-cardiac sS1 domain and intrinsic force measurements for three converter domain HCM mutant forms of the protein. **A.** Structure of homology-modeled (see Methods) human β-cardiac sS1, which contains residues 1-808 of the MyHC (dark grey) and the ELC (beige). The positions of the HCM mutations R719W (blue), R723G (red) and G741R (green) in the converter domain (white) are shown. **B.** Blowup of the converter domain shown in A. **C.** Intrinsic force measurements using a dual-beam laser trap. More than 3 independent protein preparations were made for each mutant and for WT human β-cardiac sS1, 800-900 binding events out of 6-7 individual molecules were analyzed. P=0.0067 (one-way ANOVA). The error bars show the SEM. **D.** Cumulative probability distributions from the same intrinsic force measurements shown in C.

For reasons outlined in the introduction, we expected that the converter mutations R719W, R723G and G741R (Fig. 1A,B) might cause an increase in F_intrinsic_ of the human β-cardiac sS1, and thereby explain the hyper-contractility caused by HCM mutations clinically. We used a single-molecule laser-trap *(9, 10, 12)* to measure the intrinsic forces of the 3 converter HCM mutations compared to the WT human β-cardiac sS1 (see Materials and Methods).

The force data were analyzed by multiple methods to confirm that the comparison of force-producing ability between wild type (WT) and mutants is unbiased (See Materials and Methods). Fig. 1C shows the summary of average F_intrinsic_ measurements for the 3 converter domain mutant proteins and WT. Surprisingly, compared to WT, R719W and R723G showed ~15-30% reduction in Fintrinsic, while there was no significant difference for G741R human β-cardiac sS1. An alternative way to approach the data is to look at a cumulative probability distribution of all individual intrinsic forces measured (Fig. 1D). This measures the probability of occurrence of any particular force event; for example, at a cumulative probability distribution of 0.5 the intrinsic force value was lower for R719W and R723G compared to WT, suggesting that single molecules of R719W and R723G are more probable to generate a lower intrinsic force compared to WT. Thus, these changes would contribute to either no change (G741R) or to a small hypo-contractility contribution (R719W, R723G) toward power output.

### ATPase activity was not affected by the converter domain mutations

Next we compared the actin-activated ATPase activity of recombinant sS1 carrying the converter domain mutations with WT human β-cardiac sS1 (e.g., Fig. 2A; Fig S1B-C). The actin-activated myosin ATPase activity of each mutant myosin was normalized to the WT myosin from protein preparations made simultaneously to minimize the effect of variability between protein batches (Fig. 2B). All three mutations showed no significant change in the maximal rate of actin-activated myosin ATP hydrolysis (k_cat_) (Fig. 2B, Fig. S2), and there were no significant differences in K_m_ for actin for any of the mutants (range was 20-30 µM actin). The inverse of k_cat_ defines the overall cycle time t_c_. Taken together, the t_c_ of the ATPase cycle was unchanged with the converter domain mutations. Thus, both the F_intrinsic_ and ATPase measurements for the 3 converter HCM mutations failed to reveal changes in parameters that could account for the hyper-contractility seen clinically for myosin-based HCM mutations.

**Figure 2.**
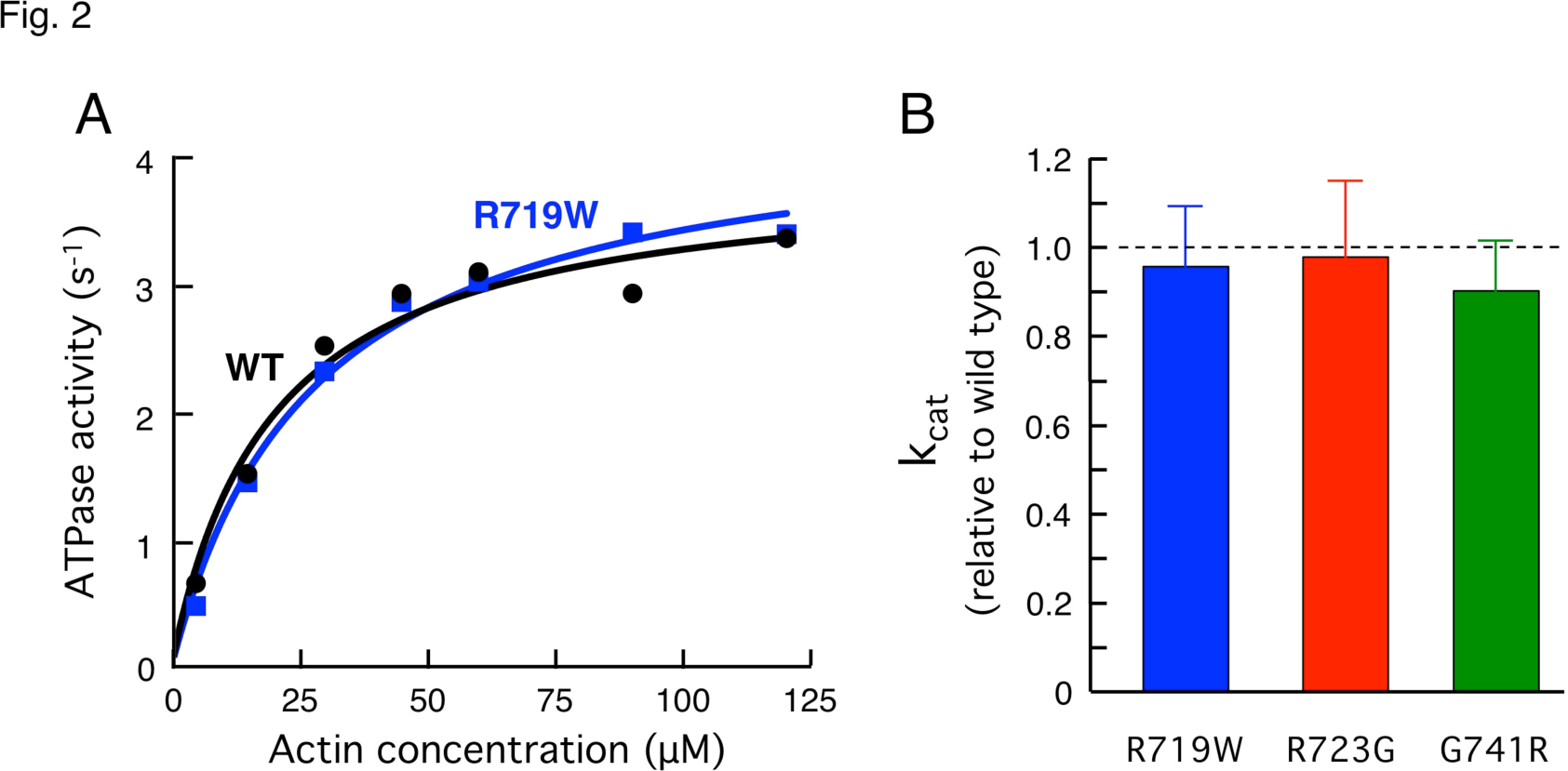
Actin-activated ATPase activity of recombinant human β-cardiac sS1 with mutations in the converter domain compared with WT human β-cardiac sS1. **A.** Representative actin-activated sS1 ATPase curves. **B.** Mutant human β-cardiac sS1 k_cat_ values relative to WT are shown (error bars: 95% confidence interval).

### Actin and regulated thin filament gliding velocities were faster in R719W and R723G human β-cardiac sS1 and unchanged in the G741R myosin

Since power output is the product of F_ensemble_ and velocity, we next assessed the mechanical property of the myosins with HCM mutations using the in vitro motility assay, where sS1 was anchored to surface-attached PDZ domain peptides that bind tightly to an 8-residue affinity clamp sequence on the C-terminus of the sS1 (see Materials and Methods). A significant increase in velocity by the mutant proteins could offset the apparent small decrease in F_ensemble_, resulting in a net increase in power output. We therefore measured the velocities of actin filaments and regulated thin filaments (RTF, consisting of actin saturated with the tropomyosin-troponin complex), gliding along surfaces coated with the various myosin preparations.

The actin/RTF filaments were tracked using the previously reported FAST program *(11)*. FAST enables tracking and analysis of hundreds of actin filaments and provides a plot of filament length-velocity relationships as well as histograms (e.g., Fig. 3A). Here we focus on 3 parameters: The top 5% of velocities for each myosin preparation (Top5%), the mean velocity (MVEL) and the percent of the filaments that are mobile (% mobile; see Materials and Methods).

**Figure 3.**
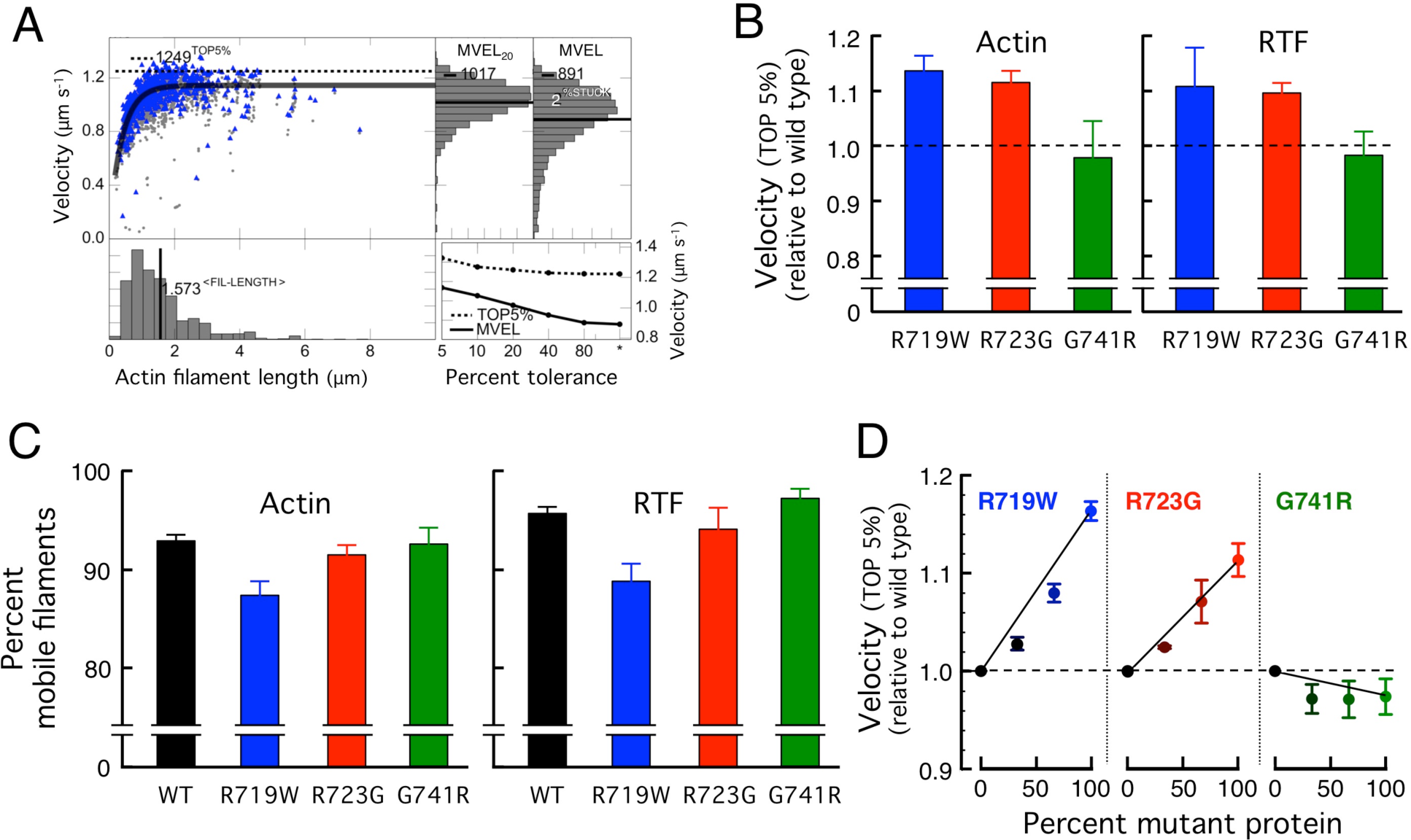
In vitro motility data for the three mutant human β-cardiac sS1 proteins compared to WT. **A**. An example of automatic analysis of an in vitro motility movie with FAST *(11)* (available for download at http://spudlab.stanford.edu). A scatterplot is shown of actin filament velocities as a function of filament length (grey, all velocity points for each filament; blue, maximum velocity of each filament, and the dashed line indicates the TOP5% of filament velocities. Also shown in the upper right are two histograms of velocities: a histogram of all filament velocities with its mean velocity (MVEL) marked as a black line, and a histogram of velocities with tolerance filtering to eliminate intermittently moving filaments with a velocity dispersion higher than 20% of their mean within a 5-frame window (MVEL_20_). In the lower right the effects of various levels of tolerance filtering on the TOP5% and the MVEL velocities are shown. **B**. The TOP5% velocities relative to WT are shown for gliding actin filaments (left) or RTFs (in 10^-^5 M Ca^2+^, right) driven by the three mutant human β-cardiac sS1 proteins. Each mutant protein was normalized against its matching WT protein prepared and assayed on the same days. Each bar graph is a mean of relative velocity (Error bars represent ± 95% confidence interval). **C**. The percent mobile filaments for the velocity measurements using WT and mutant human β-cardiac sS1 proteins, for actin (left) and RTFs (right). Data is presented as mean ± SEM. **D**. In vitro motility assays with mixtures of WT and mutant protein at varyingratios.

R719W and R723G human β-cardiac sS1 showed ~12-15% increase in Top5% gliding velocity, whereas no significant difference was observed for the G741R sS1 (Fig. 3B). When mean velocity (MVEL) was calculated, however, R719W human β-cardiac sS1 no longer showed an increase compared to WT, while R723G still retained a small ~5% increase in velocity for actin and for RTF (Fig S3).

The difference between the Top5% and MVEL levels are likely explained by the difference in the amount of stuck filaments for each mutant, which is plotted as % mobile filaments in Fig. 3C. The motility surface of the R719W mutant showed significantly more stuck filaments than WT (and hence less % mobile filaments), adding more drag force to moving filaments. While the R719W showed faster velocity when we only scored the fastest moving filaments (i.e., Top5%), which are least affected by the drag force from stuck filaments, the average velocity (i.e., MVEL) taken from the entire histogram is brought down to a similar level of WT by the presence of this drag force.

The important conclusions from the above data are that all the above changes are small and vary from slightly hypo-contractility to slightly hyper-contractility contributions, and it is not possible to explain the clinical hyper-contractility seen in HCM patients by changes in these parameters.

Note that all of the above studies were carried out with pure populations of mutant or WT proteins, whereas patients carrying these mutations are heterozygous. It has been previously reported that HCM patients with R719W and R723G mutations express ~70% of mutant protein in their cardiomyocytes *(39)*. Thus, it is possible that one would observe significant alterations in, for example, the velocity of actin filament movement if mixtures of mutant and WT proteins were examined. While we observed a dose-dependent increase in the velocity of gliding filaments as the percent of R719W and R723G mutant proteins were increased, there were no dramatic cooperative effects seen (Fig. 3D). Rather the mixtures showed velocities that were near those expected by weight-averaging the velocities of the mutant and WT proteins, with perhaps a slightly higher influence by the WT human β-cardiac sS1 for the R719W/WT mixtures and a slightly higher influence by the G741R human β-cardiac sS1 for the G741R/WT mixtures (Fig. 3D).

### The three converter HCM human β-cardiac sS1 proteins showed load-dependent velocity-change profiles that are very similar to wild type

The above velocity experiments were carried out under “unloaded” conditions, meaning that there was no additional load applied to the myosin in the in vitro motility assays. However, in cardiovascular physiology the heart is always operating under varying amount of preload and afterload, and we therefore compared the behaviors of the three converter HCM mutant human β-cardiac sS1 proteins to WT human β-cardiac sS1 using a loaded in vitro motility assay.

Thus, to the myosin-coated surface we added varying amounts of a recombinant protein that binds to actin and places a load on the myosin *(11)*. The loaded in vitro motility assay shows a relationship between filament velocity and the concentration of load molecule that relates to the force-velocity relationship of contracting muscle. As a loading molecule, we used a utrophin construct with the same 8-residue C-terminal affinity clamp tag that is at the C-terminus of the sS1 construct *(11, 12)*. At zero utrophin concentration the actin filaments glide at their maximal velocity, as there is no additional load on the myosin. As the concentration of the utrophin increases the velocity of filaments decreases. The differences in the degree change in velocity intuitively correspond to the differences in ensemble force produced by the myosin molecules on the surface; the higher the ensemble force, the less effect is seen by the utrophin load molecule. The concept of a loaded in vitro motility assay has been used in characterizing the ensemble force of motor proteins for many years *(11, 12, 40, 41)*.

Figure 4 shows the loaded motility assay applied to the 3 converter domain mutant proteins and WT, using either actin or RTF as gliding filaments. Each curve represents the compilation of 2-11 experiments from each of 2 to 5 fresh protein preparations, where a fresh preparation of WT human β-cardiac sS1 was made simultaneously with every mutant protein preparation so they could be compared under identical preparation and experimental conditions. The striking result is that, perhaps with the exception of the R723G human β-cardiac sS1 driving RTF gliding, the curves are nearly identical between the mutant proteins and WT, in keeping with the very small changes in F_intrinsic_, k_cat_ and unloaded velocities described above.

**Figure 4.**
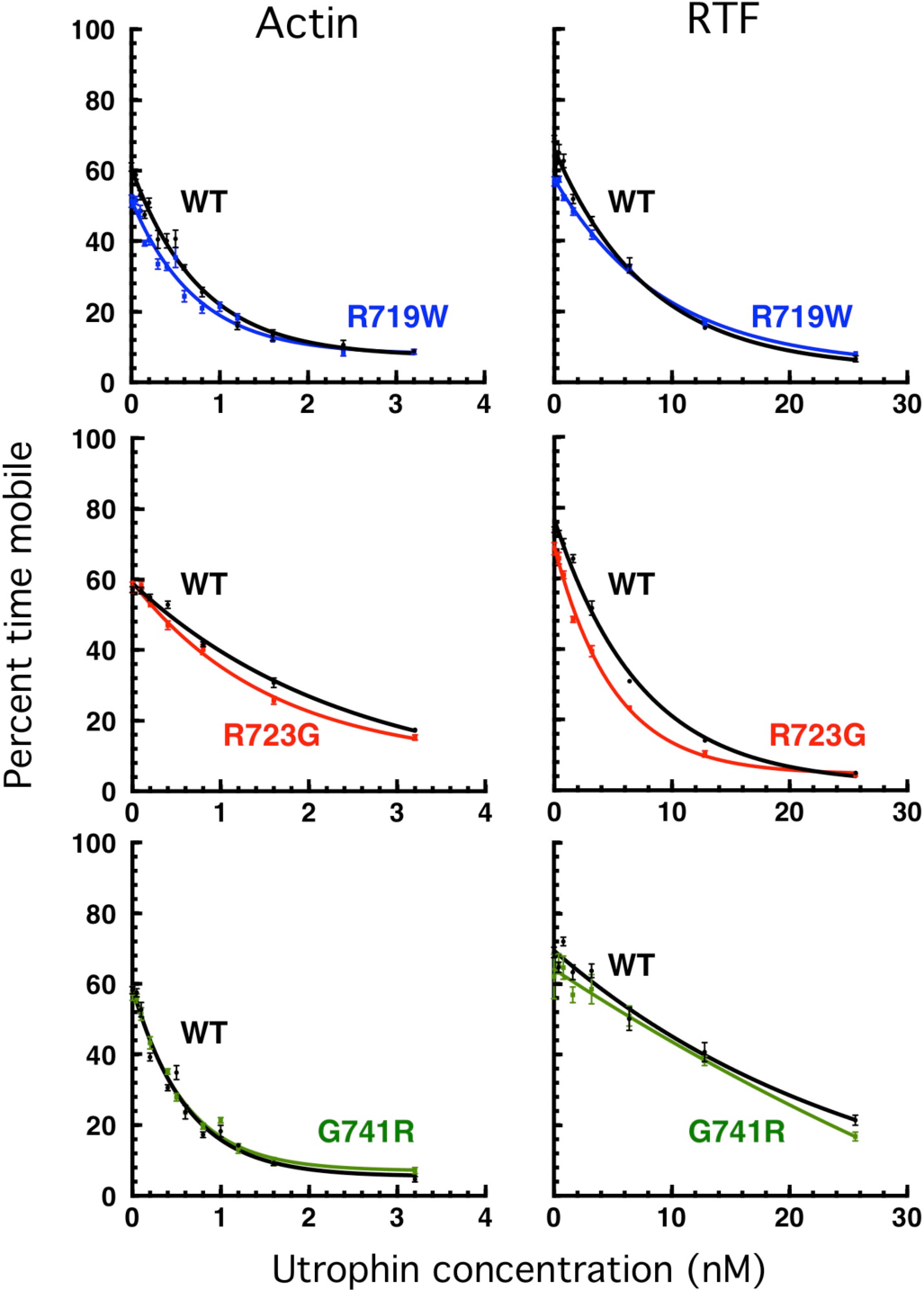
Loaded in vitro motility assays for the three mutant proteins compared to WT controls. The percent time mobile *(11)* is a measure of the relative effectiveness of the load molecule utrophin to overcome the F_ensemble_ of the sS1 on the surface *(11)* (see Materials and Methods). The effect of utrophin concentration on the velocity of actin (left) and RTFs (right, in 10^-5^ M Ca^2+^) are shown. For WT and each mutant protein, ≥5 independent protein preparations and ≥5 separate sets of curves were combined.

Thus, quite unexpectedly, the fundamental parameters for power output that we report on here for the converter mutant human β-cardiac sS1 proteins are very similar to WT human β-cardiac sS1, and we are driven to consider a different molecular basis for hyper-contractility for myosin-based HCM mutations (see Discussion).

## Discussion

The conventional view on clinical hyper-contractility resulting from HCM mutations is emphasized by the work by Ho and colleagues on patients who carry known HCM mutations but have normal left ventricular thickness and left ventricular size (so called genotype+/phenotype-individuals). These patients show a combination of diastolic dysfunction and increased ejection fraction, as measured by echocardiography with 2D tissue Doppler imaging *(7)*, suggesting that the increase in ejection fraction could not be attributed to a decrease in stroke volume, which is commonly seen in more advanced patients who manifest left ventricular hypertrophy and decrease in left ventricular chamber size. Hence, it is generally thought that HCM mutations result in gain of function.

The converter domain is the most frequently occurring domain for HCM mutations within the entire myosin molecule *(17, 18)*, and its function has important implications in force generation, load-dependent ADP release, ATP hydrolysis and Pi release. Our examination of the relevant biochemical and biophysical parameters for three key converter domain HCM mutations, R719W, R723G and G741R, showed that R719W and R723G both showed significant reduction in F_intrinsic_, and increased velocity in the in vitro motility assay, likely due to a decrease in t_s_, and downward shift in the loaded in vitro motility assay. As for the G741R mutation, there was no obvious change measured with the present experimental assays, and it appears that the mutation has neutral effect on the biochemical and biophysical functions of the molecule.

The affected parameters in the present study show even smaller magnitudes of change compared to either R403Q *(12)* or R453C human β-cardiac sS1 *(10)*, and the direction of change was for the most part opposite to our hypothesis that HCM mutations cause gain of function. It can be argued that these hypo-contractile changes could lead to left ventricular pressure and/or volume overload due to mild left ventricular systolic dysfunction, leading to compensatory hypertrophy *(42,43)*. It is also important to point out from this study that the effect of the mutation was variable between different locations and amino acid change, even within the same domain of the molecule. Hence, it is possible that the mutations selected for this study do not result in hyper-contractile changes at the sarcomere level, and there are alternative mechanisms involved in pathogenesis.

Yet, there is one pivotal parameter that has been largely neglected in studies of HCM mutations, the number of functionally accessible heads (N_a_) for interaction with actin. We have recently proposed that the majority of known myosin-based HCM mutations cause an increase in N_a_, which results in an increase in F_ensemble_, and therefore in hyper-contractility seen clinically *(44, 45)*. For reasons described below, we hypothesize that the three converter mutations studied here fall in this category.

There are many studies that demonstrate that striated muscle myosins exist in both an open state, with each head (Subfragment 1 of myosin, or S1) available for interaction with actin, and a closed state, where the heads are folded back onto their own coiled-coiled tail (Subfragment 2 of myosin, or S2) *(46-51)* (Fig. 5A). The first folded structure of this type dates back to 2001 when Wente et. al. *(52)* showed for smooth muscle myosin that the ‘blocked head’ (so-called because its actin binding face is nor accessible for binding to actin; Fig. 5B, dark grey head) interacts with the converter domain of the ‘free head’ (Fig. 5B, light grey head with white converter). Subsequently, the same folded structure has been seen in a variety of other myosins, including skeletal *(46)* and cardiac *(53)*, and Fig. 5 shows the structure of Alamo et al. *(47)*, which we homology modeled for human β-cardiac myosin (see Materials and Methods).

**Figure 5.**
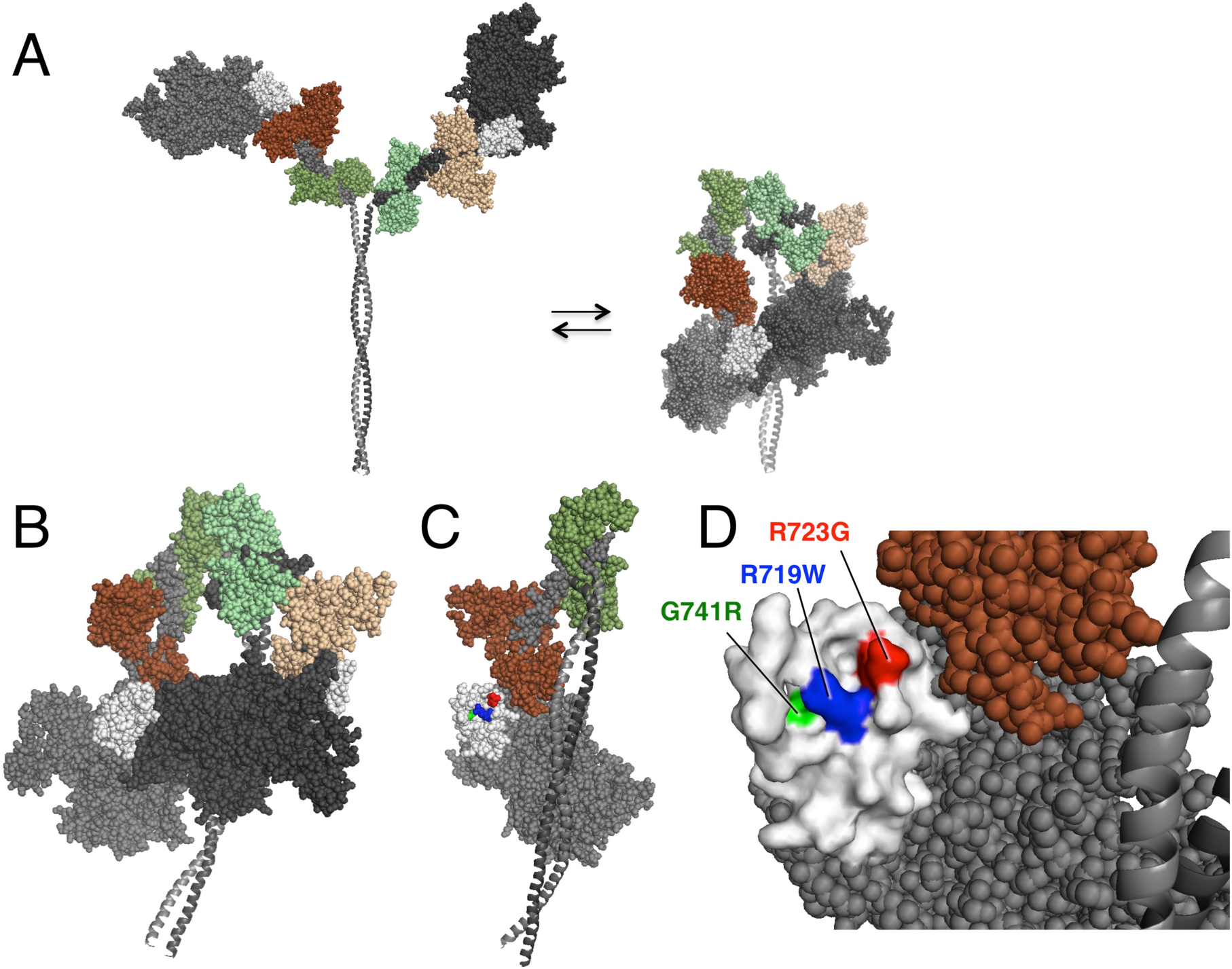
Homology-modeled structures of human β-cardiac short HMM (MyHC ending at residue 808) in the open and closed states. **A**. Illustration of the equilibrium between open and closed states. **B**. The closed state, shown enlarged and slightly rotated to the left about its vertical axis compared to that in A. The interface of the blocked head (dark grey) and the converter of the free head (white) is marked by the red line. **C**. Face-on view of the converter domain surface involved in the S1-S1 interaction. This view was obtained by rotating the molecule in B left 90° about its vertical axis and removing the blocked head from the image. **D**. Blowup of the binding interface of the converter in C, showing the positions of the three converter domain HCM mutations studied here.

What’s striking is what one observes when rotating the molecule in Fig. 5B 90° to the left about its vertical axis and removing the blocked head, so one is looking at the binding interface of the converter (Fig. 5C,D). R719 and R723 are right on the surface of the binding interface, and G741 is slightly below the surface where a change to an arginine is likely to be generally disruptive to the surface architecture. The binding interface of the blocked head is generally negatively charged *(45)*, such that R719 and R723 seem certainly important for the S1-S1 interaction shown.

As a precedent for this hypothesis, Nag et al. *(45)* have provided experimental evidence for an interaction between sS1 and the proximal region of the S2 coiled-coil tail, as well as with the C0-C2 domain of MyBP-C, both hypothesized to be occurring on the other side of the folded molecule from that shown in Fig. 5B. Nag et al. *(45)* showed that these interactions are weakened as a result of myosin HCM mutations on the myosin mesa, in a manner consistent with the structural model shown in Fig. 5.

Thus, our future work on these and other converter mutations, nearly all of which are near the S1-S1 binding interface *(45)*, will be to experimentally explore our hypothesis that the converter HCM mutations generally are weakening the S1-S1 interaction, causing an increase in N_a_ and therefore causing the hyper-contractility observed clinically.

## Material and Methods

### Myosin constructs and protein expression

Human β-cardiac myosin S1 with 3 converter domain mutations (R719W, R723G and G741R) were constructed and produced using a modified AdEasy Vector System (Qbiogene Inc.). The cloning, expression, and purification methodologies are described in detail elsewhere *(10, 12)*. Briefly, complementary DNA (cDNA) for MYH7 (human β-cardiac myosin heavy chain) and MYL3 (human ventricular ELC) were purchased from Open Biosystems (Thermo). A truncated version of MYH7 (residues 1 to 808), corresponding to a sS1, followed by a flexible GSG (Gly-Ser-Gly) linker was made with either a C-terminal enhanced green fluorescent protein (eGFP) linker (for ATPase and single-molecule optical trap measurements), or a C-terminal eight-residue (RGSIDTWV) PDZ binding peptide (for motility and ADP release experiments) and was coexpressed with human ventricular ELC containing an N-terminal FLAG tag (DYKDDDDK) and tobacco etch virus (TEV) protease site in mouse myoblast C2C12 cells (Fig. S1A). Purified fractions were stored in column buffer (10 mM imidazole, 4 mM MgCl2, 1 mM ATP, 1 mM dithiothreitol (DTT), and ~200 mM NaCl) containing 10% sucrose and were flash-frozen in liquid nitrogen before storage at −80°C. Frozen proteins exhibited similar ATPase and motility properties as compared to their fresh counterparts. Before any experiment, the myosin constructs were exchanged into the appropriate buffer conditions using Amicon centrifugal filter units (Millipore) followed by centrifugation at 350,000g for at least 10 min to remove any aggregated protein.

### Additional protein purification

#### Actin

Chicken and bovine α-cardiac actin were prepared from muscle acetone powders, using slight modifications of previously described protocols *(54)*. In addition, we used purified bovine G-actin gifted to us by MyoKardia Inc. After preparation, actin was stored in its F form in 2 mM tris (pH 8), 50 mM KCl, 0.2 mM CaCl_2_, 2 mM ATP, 2 mM MgCl_2_, 1 mM DTT, and 0.02% sodium azide. Actin was cycled from G-to F-actin freshly for each assay and used only for up to a week before being recycled again. For optical trap measurements, biotin-labeled rabbit skeletal G-actin was purchased from Cytoskeleton. F-actin was stored at 4°C, and G-actin was frozen at −80°C for future use. ATPase experiments were done with chicken skeletal actin, and motility assays were done with α-cardiac actin.

#### Gelsolin

Full-length human gelsolin was expressed and purified on the basis of previous methods *(10, 55)*. Final fractions of gelsolin were dialyzed into buffer D *(55)*, flash-frozen in liquid nitrogen, and stored at −80°C.

#### Tropomyosin

Tropomyosin was purified from bovine cardiac tissue according to the protocol of Smillie *(56)* with modifications as described by Sommese et al. *(57)*. Purified tropomyosin was dialyzed in 20 mM imidazole (pH 7.5), 300 mM KCl, and 1 mM DTT before flash-freezing and storing at −80°C.

#### Troponin

Human adult cardiac troponin subunit (TNNT_2_, TNNI_3_, and TNNC_2_) expression and purification were based on previously published methods *(12, 57-59)*. TnT, TnI, and TnC were purified and stored in storage buffer containing 20 mM imidazole (pH 7.5), 1 M KCl, 1 mM MgCl_2_, and 1 mM DTT. These proteins were then flash-frozen and stored at −80°C for future use. Troponin complexes were formed according to Szczesna et al. *(60)* with slight modifications. Components were mixed at a molar ratio of 1.3:1.3:1 (TnI/ TnT/TnC) for 1 hr on ice. Complexes were then dialyzed at 4°C in six sequential steps into complex buffer [20 mM imidazole (pH 7.5), 2 mM MgCl_2_, and 1 mM DTT] containing 0.7, 0.5, 0.3, 0.1, and 0.01 M KCl twice, for 6 to 12 hours each. Complexes were flash-frozen in complex buffer before storing them at −80°C. The cDNAs for human adult cardiac TnI, TnC, and TnT in carbenicillin-selective pET-3d plasmids were obtained from J. Potter (University of Miami).

RTF complex formation: For all experiments involving the RTFs, RTFs were formed by mixing excess tropomyosin and troponin complex to actin on ice and then incubating for at least 1 hour prior to use. The final molar ratio was 7:2:2 of actin/tropomyosin/troponin for all experiments.

#### Utrophin

Mouse utrophin with an eight-residue (RGSIDTWV) C-terminal PDZ binding peptide was expressed in bacterial cells as previously published *(11)*. The purified protein was concentrated and dialyzed overnight against 150 mM NaCl, 25 mM tris, and 1 mM DTT (pH 8.0) at 4°C before flash-freezing in liquid nitrogen and storing at −80°C.

#### PDZ18

The SNAP-PDZ18 fusion construct was expressed in bacterial cells as previously described *(11)*. Eluted protein was concentrated and the buffer was exchanged to 150 mM NaCl and 25 mM tris (pH 8.0). The purified protein was flash-frozen in liquid nitrogen and stored at −80°C.

### Actin-activated ATPase assay

For actin-activated ATPase, gelsolin was added to actin at a ratio of 1:500. Gelsolin at this concentra-tion was used to decrease the viscosity of the actin and thereby decrease pipetting error and allow higher concentrations of actin to be used. Actin-activated ATPase assays were then performed as previously described using a colorimetric readout *(61)*. Briefly, sS1 was diluted to a final concentration of 0.07 ~ 0.15 µM (with 4 times as much for the basal myosin ATPase control in the absence of actin to amplify the signal) with 2 mM ATP and actin at concentrations ranging from 0 to 125 uM.. The final buffer conditions were 10 mM imidazole (pH 7.5), 5 mM KCl, 4 mM MgCl_2_, and 1 mM DTT. The reaction was performed at 23°C with shaking in a Thermo Scientific Multiskan GO, and 4 to 5 time points were taken for each concentration. The sS1 activity was linear over the time period of the assay, and hence, an ATP-regenerating system was not necessary. Basal activity (<0.2 s^−1^) was subtracted to get actin-activated ATPase activity. The Michaelis-Menten equation was fit to the data to determine the maximal activity (k_cat_) and the associated actin constant for myosin (K_m_) using Prism 6 (Graphpad Software Inc.). All of the experiments were done using sS1 with a FLAG tag on the ELC as previously described *(10, 12)*, except for R723G which was assayed with and without the FLAG tag to check whether the presence of the FLAG tag had any effect on the k_cat_ of the sS1. The FLAG tag was removed during the purification process by adding TEV protease to cleave the myosin that was bound to the FLAG-resin. The sS1 was then purified by HPLC. No significant difference was seen between the sS1 with and without the FLAG tag (Fig. S2).

### Unloaded and loaded in vitro motility

The basic method followed our previously described motility assay *(62)*, with some modifications. Coverslips (VWR micro cover glass) were coated with a mixture of 0.2% nitrocellulose (Ernest Fullam Inc.) and 0.2% collodion (Electron Microscopy Sciences) dissolved in amyl acetate (Sigma) and air-dried for a few hours before use. A permanent double-sided tape (Scotch) was used to construct four channels in each slide (Gold Seal) and four different experiments were performed on the same slide. In general, both WT and mutant protein(s) were studied on the same slide to minimize variability in slide preparation for measurement of unloaded motility. For loaded in vitro motility assays, 2-3 slides were used under 8-12 different concentrations of utrophin to obtain a full curve with each run of an experiment.

Ten-fold molar excess of F-actin was added in the presence of 4 mM ATP, incubated for 10 min, and sedimented at 350,000g for 20 min (termed “dead-heading”) to reduce the number of partially inactivated myosin heads in S1 preparations. MgCl_2_ was added to 50 mM (to form F-actin paracrystals) and incubated for 20 min (with mixing by pipetting at 10 min), and the mixture was re-sedimented at 350,000g for 30 min to eliminate sS1 that remained bound to actin and residual actin in the supernatant. The supernatant was collected and the sS1 concentration was measured using the Bradford reagent (Bio-Rad). A mock clean-up procedure containing storage buffer, actin and MgCl2 without sS1 was also performed simultaneously and was used as the blank for more accurate concentration determination of sS1. The quality of the sS1 clean-up was assessed by the percentage of stuck filaments under unloaded conditions. We repeated the sS1 clean-up procedure until the percentage of stuck filaments dropped below 10%. Before any experiments, dead-headed sS1 was diluted in 10% ABBSA {assay buffer [AB; 25 mM imidazole (pH 7.5), 25 mM KCl, 4 mM MgCl_2_, 1 mM EGTA, and 1 mM DTT] with bovine serum albumin (BSA; 0.1 mg/ml) diluted in AB}, unless otherwise stated.

For motility experiments, reagents were sequentially flowed through the channels in the following order: (i) 10 µl of 3µM SNAP-PDZ18 diluted in AB and incubated for 2 min; (ii) 30 ml of ABBSA to block the surface from nonspecific attachments and incubated for 2 min; (iii) 10 µl of a mixture of eight-residue (RGSIDTWV)–tagged human cardiac S1 (~0.05 to 0.1 mg/ml for actin motility and 0.1 to 0.2 mg/ml for RTF motility) and utrophin at desired concentrations and incubated for 5 min (before mixing sS1 and utrophin, sS1 and utrophin dilutions were prepared in 10% ABBSA; for unloaded motility, utrophin was skipped in this step); (iv) 30 µl of ABBSA to wash any unattached proteins; and (v) finally, 10 µl of the GO solution {1 to 5 nM tetramethylrhodamine (TMR)-phalloidin (Invitrogen)–labeled bovine actin, 2 mM ATP (Calbiochem), an oxygen-scavenging system [0.2% glucose, glucose oxidase (0.11 mg/ml; Calbiochem), and catalase (0.018 mg/ml; Calbiochem)], and an ATP regeneration system [1 mM phosphocreatine (Calbiochem) and creatine phosphokinase (0.1 mg/ml; Calbiochem)]} in ABBSA.

For RTF motility experiments, higher salt concentration was necessary to avoid aggregation of RTF in solution. We also attached the RTF with myosin on the surface in the absence of ATP, and then myosin was activated by the addition of saturating concentration of ATP. Thus, after attaching the myosin at step (iii), the motility surface was washed with AB that contained 100 mM KCl, followed by addition of RTF dilution in ABBSA containing 100 mM KCl, 5 mM CaCl2, 100 nM excess tropomyosin and troponin complex and 1-5 nM TMR-phalloidin-labeled RTF. After 5 min of RTF binding, the surface was washed with ABBSA with 25 mM KCl to bring the salt concentration down. Final GO solution included AB (25 mM KCl), 5 mM CaCl2, 100 nM excess tropomyosin/troponin complex and oxygen-scavenging system as described above, and thus the motility was done at the same salt concentration as the actin experiment. The RTF mixture was made at least 1 hr before the motility was measured by mixing TMR-phalloidin-labeled bovine actin/bovine tropomyosin/human troponin complex in a 7:2:2 ratio (3.5 mM actin, 1 mM tropomyosin, 1 mM troponin and 1 mg/ml BSA).

For all experiments, movies were obtained at 23°C, at a frame rate of 1 Hz using a Nikon Ti-E inverted microscope with an Andor iXon+EMCCD camera model DU885. All experiments were repeated with at least 4 different fresh protein preparations. At each condition, at least 3 different movies with duration of 30 s were recorded. Filament tracking and analysis of movies, both under unloaded and loaded conditions, were performed by a recently published method, FAST (Fast Automated Spud Trekker), as described by Aksel et al. *(11)*

### Single-molecule optical trap measurements

The dual-beam optical trap instrumentation is described in detail elsewhere *(10,12)*. The experimental condition is similar to the one described in the motility assay. Here, the myosin construct had an eGFP tag at the C-terminal end for surface attachment through binding with anti–green fluorescent protein antibody. Dead-heads of purified protein were eliminated as described before *(10)*. All the experiments were performed at 23°C. The nitrocellulose-coated glass surface of sample chamber was also coated with 1.5 µm-diameter silica beads that acted as platforms. The sequence of steps for preparing the chamber is: (i) anti-green fluorescent protein antibody (~0.01 mg/ml) (Abcam) was flowed through the chamber, (ii) AB buffer containing 1 mg/ml BSA (ABBSA buffer) was flowed to block the exposed surface, (iii) ~50-200 pM of human β-cardiac sS1 was to sparsely coat the surface with myosin, (iv) the chamber was washed with AB buffer, and finally (v) ABBSA buffer containing 250-500 pM ATP, TMR-phalloidin–labeled biotin-actin filaments, neutravidin-coated polystyrene beads (Polysciences), and the oxygen-scavenging and ATP regeneration systems described above was flowed through the chamber. The chamber was sealed with vacuum grease to stop evaporation of the solution. Neutravidin-coated polystyrene beads (1-µm diameter) bound to each end of a TMR-phalloidin-and biotin-labeled actin filament were trapped in two different laser beams. The bead-actin-bead assembly is known as a dumbbell, which was stretched to remove compliance in the actin filament and brought close to the bead pedestal on the surface for interaction with myosin. A detailed isometric force measurement procedure is described elsewhere *(10, 12)*. A trap stiffness of ~0.1 pN/nm was used.

Individual force events were collected from several single molecules of multiple protein purifications. The number of total force events from individual molecules on average was 30 to 400. We have always observed that for all of our sS1 constructs, the force distribution is accompanied with a long tail *(10, 12)*. This phenomenon is commonly reported in the single-molecule force measurements of myosin *(63-65)*, although the exact reason is not known. It is important to take account of the smaller population of higher force events in the analysis. Hence, we chose a double Gaussian function to fit the force histogram data and the major first peak of the fit yielded the intrinsic force of an individual molecule reported here. Such intrinsic force values of multiple molecules were used to calculate the mean force value. Additionally, we have used cumulative probability distribution analysis *(12)* to compare the force-producing ability of different myosins. This method of analysis takes account of all the events to generate the function. The probability distribution at any particular force value is calculated by the adding the number of events up to that force value divided by the total number of events of all force values. Such a distribution starts from a value close to zero at the lowest measurable force to 1 at the maximum force. If the force producing ability between two proteins is different, the probability value at any force value will be different.

### Development of human β-cardiac myosin protein models

We developed human β-cardiac myosin S1 models based on the known motor domain structural data to best represent the human β-cardiac myosin, as described in Homburger et al. *(17)*. In brief, we retrieved the protein sequence of human β-cardiac myosin and the human cardiac light chains from UNIPROT database *(66)*: myosin heavy chain motor domain (MYH7) - P12883, myosin essential light chain (MLC1) - P08590, and myosin regulatory light chain (MLC2) - P10916. We used a multi-template homology modeling approach to build the structural coordinates of MYH7 (residues 1-840), MCL1 (residues 1-195), MCL2 (residues 1-166) and S2 (residues 841-1280). We obtained the three dimensional structural model of S1 in the pre-and post-stroke states by integrating the known structural data from solved crystal structures, as described *(17)*. Fig. 5 shows a homology-modeled folded-back structure of human β-cardiac myosin from the 3D-reconstructed images of tarantula skeletal muscle thick filaments by Alamo et al. *(47)*.

The S2 region is a long coiled-coil structure; hence we used the template from the Myosinome database *(67)*. Modeling was done using the MODELLER package. Visualizations were performed using PyMOL version 1.7.4 (www.pymol.org).

## Acknowledgments

The authors thank Suman Nag and Ruth Sommese for preparation of the tropomyosin and troponin complex, and Allan Borrayo for his technical help of maintaining virus and cell stocks. This project was funded by NIH Grants GM33289 and HL117138 to J.A.S. M.K. was a recipient of the NRSA Postdoctoral Fellowship F32HL124883. Myosin and actin preparations were performed by M.K. with help from S.S. and K.M.R. Steady state ATPase and in vitro motility assays were performed by M.K. Intrinsic force measurements were done by S.S.S. M.K. and J.A.S. wrote the manuscript. M.K., S.S.S., K.M.R. and J.A.S. contributed to data analysis/interpretation and editing of the manuscript. A complete version of the FAST program (*11*) for motility analysis is available for downloading on our website (http://spudlab.stanford.edu).

J.A.S. is a founder of and owns shares in Cytokinetics, Inc. and MyoKardia, Inc., biotech companies that are developing therapeutics that target the sarcomere.

## References

1. C. Semsarian, J. Ingles, M. S. Maron, B. J. Maron, New perspectives on the prevalence of hypertrophic cardiomyopathy, Journal of the American College of Cardiology 65, 1249–1254 (2015).

2. P. M. Elliott, J. R. Gimeno, R. Thaman, J. Shah, D. Ward, S. Dickie, M. T. Tome Esteban, W. J. McKenna, Historical trends in reported survival rates in patients with hypertrophic cardiomyopathy, Heart 92, 785–791 (2006).

3. A. A. Geisterfer-Lowrance, S. Kass, G. Tanigawa, H. P. Vosberg, W. McKenna, C. E. Seidman,J.G. Seidman, A molecular basis for familial hypertrophic cardiomyopathy: a beta cardiac myosin heavy chain gene missense mutation, Cell 62, 999–1006 (1990).

4. L. Wang, J. G. Seidman, C. E. Seidman, Narrative review: harnessing molecular genetics for the diagnosis and management of hypertrophic cardiomyopathy, Annals of Internal Medicine 152, 513–520 (2010).

5. B. J. Maron, M. S. Maron, C. Semsarian, Genetics of Hypertrophic Cardiomyopathy After 20 Years, Journal of the American College of Cardiology 60, 705–715 (2012).

6. J. G. Seidman, C. Seidman, The genetic basis for cardiomyopathy: from mutation identification to mechanistic paradigms, Cell 104, 557–567 (2001).

7. C. Y. Ho, N. K. Sweitzer, B. McDonough, B. J. Maron, S. A. Casey, J. G. Seidman, C. E. Seidman,S. D. Solomon, Assessment of diastolic function with Doppler tissue imaging to predict genotype in preclinical hypertrophic cardiomyopathy, Circulation 105, 2992–2997 (2002).

8. C. E. Seidman, J. G. Seidman, The Metabolic and Molecular Bases of Inherited Disease C. R. Scriver, A. L. Beaudet, D. Valle, W. S. Sly, B. Childs, K. W. Kinzler, B. vogelstein, Eds. (McGraw-Hill Professional, 2000), pp. 5433–5452.

9. J. Sung, S. Sivaramakrishnan, A. R. Dunn, J. A. Spudich, 14 - Single-Molecule Dual-Beam Optical Trap Analysis of Protein Structure and Function (Elsevier Inc., ed. 1, 2010), pp. 321–375.

10. R. F. Sommese, J. Sung, S. Nag, S. Sutton, J. C. Deacon, E. Choe, L. A. Leinwand, K. Ruppel, J.A. Spudich , Molecular consequences of the R453C hypertrophic cardiomyopathy mutation on human β-cardiac myosin motor function, Proc. Natl. Acad. Sci. U.S.A. 110, 12607–12612 (2013).

11. T. Aksel, E. Choe Yu, S. Sutton, K. M. Ruppel, J. A. Spudich, Ensemble Force Changes that Result from Human Cardiac Myosin Mutations and a Small-Molecule Effector, Cell Rep (2015), doi:10.1016/j.celrep.2015.04.006.

12. S. Nag, R. F. Sommese, Z. Ujfalusi, A. Combs, S. Langer, S. Sutton, L. A. Leinwand, M. A. Geeves, K. M. Ruppel, J. A. Spudich, Contractility parameters of human β-cardiac myosin with the hypertrophic cardiomyopathy mutation R403Q show loss of motor function, Science Advances 1, e1500511–e1500511 (2015).

13. J. A. Spudich, Hypertrophic and Dilated Cardiomyopathy: Four Decades of Basic Research on Muscle Lead to Potential Therapeutic Approaches to These Devastating Genetic Diseases, Biophysj 106, 1236–1249 (2014).

14. E. P. Debold, J. P. Schmitt, J. B. Patlak, S. E. Beck, J. R. Moore, J. G. Seidman, C. Seidman, D.M. Warshaw, Hypertrophic and dilated cardiomyopathy mutations differentially affect the molecular force generation of mouse α-cardiac myosin in the laser trap assay, Am. J. Physiol. Heart Circ. Physiol. 293, H284–H291 (2007).

15. M. J. Tyska, E. Hayes, M. Giewat, C. E. Seidman, J. G. Seidman, D. M. Warshaw, Single-molecule mechanics of R403Q cardiac myosin isolated from the mouse model of familial hypertrophic cardiomyopathy, Circulation Research86, 737–744 (2000).

16. S. Lowey, L. M. Lesko, A. S. Rovner, A. R. Hodges, S. L. White, R. B. Low, M. Rincon, J. Gulick, J. Robbins, Functional effects of the hypertrophic cardiomyopathy R403Q mutation are different in an α-or β-myosin heavy chain backbone, J. Biol. Chem. 283, 20579–20589 (2008).

17. J. R. Homburger, E. M. Green, C. Caleshu, M. S. Sunitha, R. E. Taylor, K. M. Ruppel, R. P. R. Metpally, S. D. Colan, M. Michels, S. M. Day, I. Olivotto, C. D. Bustamante, F. E. Dewey, C. Y. Ho, J. A. Spudich, E. A. Ashley, Multidimensional structure-function relationships in human β-cardiac myosin from population-scale genetic variation, Proceedings of the National Academy of Sciences, 201606950 (2016).

18. M. Colegrave, M. Peckham, Structural Implications of β-Cardiac Myosin Heavy Chain Mutations in Human Disease, Anat. Rec. 297, 1670–1680 (2014).

19. D. García-Giustiniani, M. Arad, M. Ortíz-Genga, R. Barriales-Villa, X. Fernández,I. Rodríguez-García, A. Mazzanti, E. Veira, E. Maneiro, P. Rebolo, I. Lesende, L. Cazón, D. Freimark, J. R. Gimeno-Blanes, C. Seidman, J. Seidman, W. McKenna, L. Monserrat, Phenotype and prognostic correlations of the converter region mutations affecting the β myosin heavy chain, Heart 101, 1047–1053 (2015).

20. J. Köhler, G. Winkler, I. Schulte, T. Scholz, W. McKenna, B. Brenner, T. Kraft,Mutation of the myosin converter domain alters cross-bridge elasticity, Proc. Natl. Acad. Sci. U.S.A. 99, 3557–3562 (2002).

21. K. P. Littlefield, D. M. Swank, B. M. Sanchez, A. F. Knowles, D. M. Warshaw, S. I. Bernstein, The converter domain modulates kinetic properties of Drosophila myosin, Am. J. Physiol., Cell Physiol. 284, C1031–8 (2003).

22. W. A. Kronert, G. C. Melkani, A. Melkani, S. I. Bernstein, Mutating the Converter–Relay Interface of Drosophila Myosin Perturbs ATPase Activity, Actin Motility, Myofibril Stability and Flight Ability, Journal of Molecular Biology 398, 625–632 (2010).

23. D. M. Swank, A. F. Knowles, J. A. Suggs, F. Sarsoza, A. Lee, D. W. Maughan, S. I. Bernstein, The myosin converter domain modulates muscle performance, Nat. Cell Biol. 4, 312–316 (2002).

24. S. Ramanath, Q. Wang, S. I. Bernstein, D. M. Swank, Disrupting the myosin converter-relay interface impairs Drosophila indirect flight muscle performance, Biophysical Journal 101, 1114–1122 (2011).

25. R. Anan, G. Greve, L. Thierfelder, H. Watkins, W. J. McKenna, S. Solomon, C. Vecchio, H. Shono, S. Nakao, H. Tanaka, A. Mares, J. A. Towbin, P. Spirito, R. Roberts, J. G. Seidman, C. E. Seidman, Prognostic implications of novel beta cardiac myosin heavy chain gene mutations that cause familial hypertrophic cardiomyopathy, J. Clin. Invest. 93, 280–285 (1994).

26. M. Enjuto, A. Francino, F. Navarro-LopEz, D. Viles, J. C. Paré, A. M. Ballesta, Malignant hypertrophic cardiomyopathy caused by the Arg723Gly mutation in beta-myosin heavy chain gene, Journal of Molecular and Cellular Cardiology 32, 2307–2313 (2000).

27. L. Fananapazir, M. C. Dalakas, F. Cyran, G. Cohn, N. D. Epstein, Missense mutations in the beta-myosin heavy-chain gene cause central core disease in hypertrophic cardiomyopathy, Proc Natl Acad Sci USA 90, 3993–3997 (1993).

28. B. Seebohm, F. Matinmehr, J. KOhler, A. Francino, F. Navarro-LopEz, A. Perrot, C. Ozcelik, W. J. McKenna, B. Brenner, T. Kraft, Cardiomyopathy Mutations Reveal Variable Region of Myosin Converter as Major Element of Cross-Bridge Compliance, Biophysj 97, 806–824 (2009).

29. P. Teekakirikul, S. Eminaga, O. Toka, R. Alcalai, L. Wang, H. Wakimoto, M. Nayor, T. Konno, J. M. Gorham, C. M. Wolf, J. B. Kim, J. P. Schmitt, J. D. Molkentin, R. A. Norris, A. M. Tager, S. R. Hoffman, R. R. Markwald, C. E. Seidman, J. G. Seidman, Cardiac fibrosis in mice with hypertrophic cardiomyopathy is mediated by non-myocyte proliferation and requires Tgf-β, J. Clin. Invest. 120, 3520–3529 (2010).

30. S. E. Kirschner, E. Becker, M. Antognozzi, H.-P. Kubis, A. Francino, F. Navarro-Lopez, N. Bit-Avragim, A. Perrot, M. M. Mirrakhimov, K.-J. Osterziel, W. J. McKenna, B. Brenner, T. Kraft, Hypertrophic cardiomyopathy-related beta-myosin mutations cause highly variable calcium sensitivity with functional imbalances among individual muscle cells, Am. J. Physiol. Heart Circ. Physiol. 288, H1242–51 (2005).

31. T. Kraft, E. R. Paalberends, N. M. Boontje, S. Tripathi, A. Brandis, J. Montag, J. L. Hodgkinson, A. Francino, F. Navarro-Lopez, B. Brenner, G. J. M. Stienen, J. van der Velden, Familial Hypertrophic Cardiomyopathy: Functional effects of myosin mutation R723G in cardiomyocytes, Journal of Molecular and Cellular Cardiology, 1–31 (2013).

32. E. B. Lankford, N. D. Epstein, L. Fananapazir, H. L. Sweeney, Abnormal contractile properties of muscle fibers expressing beta-myosin heavy chain gene mutations in patients with hypertrophic cardiomyopathy, J. Clin. Invest. 95, 1409 (1995).

33. G. Miller, J. Maycock, E. White, M. Peckham, S. Calaghan, Heterologous expression of wild-type and mutant β-cardiac myosin changes the contractile kinetics of cultured mouse myotubes, The Journal of Physiology 548, 167–174 (2003).

34. W. A. Kronert, G. C. Melkani, A. Melkani, S. I. Bernstein, Alternative Relay and Converter Domains Tune Native Muscle Myosin Isoform Function in Drosophila, Journal of Molecular Biology 416, 543–557 (2012).

35. A. G. Bick, J. Flannick, K. Ito, S. Cheng, R. S. Vasan, M. G. Parfenov, D. S. Herman, S. R. DePalma, N. Gupta, S. B. Gabriel, B. H. Funke, H. L. Rehm, E. J. Benjamin, J. Aragam, H. A. Taylor Jr, E. R. Fox, C. Newton-Cheh, S. Kathiresan, C. J. O’Donnell, J. G. Wilson, D. M. Altshuler, J. N. Hirschhorn, J. G. Seidman, C. Seidman, Burden of Rare Sarcomere Gene Variants in the Framingham and Jackson Heart Study Cohorts, The American Journal of Human Genetics 91, 513–519 (2012).

36. Y. Y. Toyoshima, S. J. Kron, E. M. McNally, K. R. Niebling, C. Toyoshima, J. A. Spudich, Myosin subfragment-1 is sufficient to move actin filaments in vitro, Nature 328, 536–539 (1987).

37. T. Q. Uyeda, S. J. Kron, J. A. Spudich, Myosin step size: estimation from slow sliding movement of actin over low densities of heavy meromyosin, Journal of Molecular Biology 214, 699–710 (1990).

38. T. Q. Uyeda, P. D. Abramson, J. A. Spudich, The neck region of the myosin motor domain acts as a lever arm to generate movement, Proc. Natl. Acad. Sci. U.S.A. 93, 4459–4464 (1996).

39. B. Brenner, B. Seebohm, S. Tripathi, J. Montag, T. Kraft, Familial hypertrophic cardiomyopathy: functional variance among individual cardiomyocytes as a trigger of FHC-phenotype development, Front Physiol 5, 392 (2014).

40. M. J. Greenberg, J. R. Moore, The molecular basis of frictional loads in the in vitro motility assay with applications to the study of the loaded mechanochemistry of molecular motors, Cytoskeleton 67, 273–285 (2010).

41. M. J. Greenberg, K. Kazmierczak, D. Szczesna-Cordary, J. R. Moore, Cardiomyopathy-linked myosin regulatory light chain mutations disrupt myosin strain-dependent biochemistry, Proc. Natl. Acad. Sci. U.S.A. 107, 17403–17408 (2010).

42. D. L. Mann, Mechanisms and Models in Heart Failure: The Biomechanical Model and Beyond, Circulation 111, 2837–2849 (2005).

43. B. C. Bernardo, K. L. Weeks, L. Pretorius, J. R. McMullen,Molecular distinction between physiological and pathological cardiac hypertrophy: Experimental findings and therapeutic strategies, Pharmacology and Therapeutics 128, 191–227 (2010).

44. J. A. Spudich, The myosin mesa and a possible unifying hypothesis for the molecular basis of human hypertrophic cardiomyopathy, Biochem. Soc. Trans. 43, 64–72 (2015).

45. S. Nag, D. Trivedi, S. Sarkar, S. Sutton, K. M. Ruppel, J. A. Spudich, Beyond the myosin mesa: a potential unifying hypothesis on the underlying molecular basis of hyper-contractility caused by a majority of HCM mutations, Submitted.

46. J. L. Woodhead, F.-Q. Zhao, R. Craig, E. H. Egelman, L. Alamo, R. Padrón, Atomic model of a myosin filament in the relaxed state, Nature 436, 1195–1199 (2005).

47. L. Alamo, W. Wriggers, A. Pinto, F. Bártoli, L. Salazar, F.-Q. Zhao, R. Craig, R. Padrón, Three-dimensional reconstruction of tarantula myosin filaments suggests how phosphorylation may regulate myosin activity, Journal of Molecular Biology 384, 780–797 (2008).

48. L. Alamo, X. E. Li, L. M. Espinoza-Fonseca, A. Pinto, D. D. Thomas, W. Lehman, R. Padrón, Tarantula myosin free head regulatory light chain phosphorylation stiffens N-terminal extension, releasing it and blocking its docking back, Mol Biosyst 11, 2180–2189 (2015).

49. L. Alamo, D. Qi, W. Wriggers, A. Pinto, J. Zhu, A. Bilbao, R. E. Gillilan, S. Hu, R. Padrón, Conserved Intramolecular Interactions Maintain Myosin Interacting-Heads Motifs Explaining Tarantula Muscle Super-Relaxed State Structural Basis, Journal of Molecular Biology 428, 1142–1164 (2016).

50. N. Naber, R. Cooke, E. Pate, Slow myosin ATP turnover in the super-relaxed state in tarantula muscle, Journal of Molecular Biology 411, 943–950 (2011).

51. P. Hooijman, M. A. Stewart, R. Cooke,A new state of cardiac myosin with very slow ATP turnover: a potential cardioprotective mechanism in the heart, Biophysical Journal 100, 1969–1976 (2011).

52. T. Wendt, D. Taylor, K. M. Trybus, K. Taylor, Three-dimensional image reconstruction of dephosphorylated smooth muscle heavy meromyosin reveals asymmetry in the interaction between myosin heads and placement of subfragment 2, Proc. Natl. Acad. Sci. U.S.A. 98, 4361–4366 (2001).

53. M. E. Zoghbi, J. L. Woodhead, R. L. Moss, R. Craig, Three-dimensional structure of vertebrate cardiac muscle myosin filaments, Proceedings of the National Academy of Sciences 105, 2386–2390 (2008).

54. J. D. Pardee, J. A. Spudich, Purification of muscle actin, Meth. Enzymol. 85 Pt B, 164–181 (1982).

55. M. Way, J. Gooch, B. Pope, A. G. Weeds, Expression of human plasma gelsolin in Escherichia coli and dissection of actin binding sites by segmental deletion mutagenesis, The Journal of Cell Biology 109, 593–605 (1989).

56. L. B. Smillie, Preparation and identification of alpha-and beta-tropomyosins, Meth. Enzymol. 85 Pt B, 234–241 (1982).

57. R. F. Sommese, S. Nag, S. Sutton, S. M. Miller, J. A. Spudich, K. M. Ruppel, A. Kimura, Ed. Effects of Troponin T Cardiomyopathy Mutations on the Calcium Sensitivity of the Regulated Thin Filament and the Actomyosin Cross-Bridge Kinetics of Human β-Cardiac Myosin,PLoS ONE 8, e83403 (2013).

58. Z. Sheng, B. S. Pan, T. E. Miller, J. D. Potter, Isolation, expression, and mutation of a rabbit skeletal muscle cDNA clone for troponin I.The role of the NH2 terminus of fast skeletal muscle troponin I in its biological activity, J. Biol. Chem. 267, 25407–25413 (1992).

59. B. S. Pan, J. D. Potter, Two genetically expressed troponin T fragments representing alpha and beta isoforms exhibit functional differences, J. Biol. Chem. 267, 23052–23056 (1992).

60. D. Szczesna, G. Guzman, T. Miller, J. Zhao, K. Farokhi, H. Ellemberger, J. D. Potter, The role of the four Ca2+ binding sites of troponin C in the regulation of skeletal muscle contraction, J. Biol. Chem. 271, 8381–8386 (1996).

61. K. M. Trybus,Biochemical Studies of Myosin, Methods 22, 327–335 (2000).

62. S. J. Kron, Y. Y. Toyoshima, T. Q. Uyeda, J. A. Spudich, Assays for actin sliding movement over myosin-coated surfaces, Meth. Enzymol. 196, 399–416 (1991).

63. M. W. Allersma, F. Gittes, M. J. deCastro, R. J. Stewart, C. F. Schmidt, Two-dimensional tracking of ncd motility by back focal plane interferometry, Biophysj 74, 1074–1085 (1998).

64. R. M. Simmons, J. T. Finer, S. Chu, J. A. Spudich, Quantitative measurements of force and displacement using an optical trap, Biophysj 70, 1813–1822 (1996).

65. Y. Takagi, E. E. Homsher, Y. E. Goldman, H. Shuman, Force Generation in Single Conventional Actomyosin Complexes under High Dynamic Load, Biophysj 90, 1295–1307 (2006).

66. UniProt Consortium, Update on activities at the Universal Protein Resource (UniProt) in 2013, Nucleic Acids Research 41, D43–7 (2013).

67. D. P. Syamaladevi, M. S. Sunitha, S. Kalaimathy, C. C. Reddy, M. Iftekhar, S. N. Pasha, R. Sowdhamini,Myosinome: a database of myosins from select eukaryotic genomes to facilitate analysis of sequence-structure-function relationships, Bioinform Biol Insights 6, 247–254 (2012).

